# Insights into the genetic epidemiology of Crohn’s and rare diseases in the Ashkenazi Jewish population

**DOI:** 10.1101/077180

**Authors:** Manuel A. Rivas, Jukka Koskela, Hailiang Huang, Christine Stevens, Brandon E. Avila, Talin Haritunians, Benjamin M. Neale, Mitja Kurki, Andrea Ganna, Daniel Graham, Benjamin Glaser, Inga Peter, Gil Atzmon, Nir Barzilai, Adam P. Levine, Elena Schiff, Nikolas Pontikos, Ben Weisburd, Konrad J. Karczewski, Eric V. Minikel, Britt-Sabina Petersen, Laurent Beaugerie, Philippe Seksik, Jacques Cosnes, Stefan Schreiber, Bernd Bokemeyer, Johannes Bethge, NIDDK IBD Genetics consortium, T2D-GENES consortium, Graham Heap, Tariq Ahmad, Vincent Plagnol, Anthony W. Segal, Stephan Targan, Dan Turner, Paivi Saavalainen, Martti Farkkila, Kimmo Kontula, Matti Pirinen, Aarno Palotie, Steven R. Brant, Richard H. Duerr, Mark S. Silverberg, John D. Rioux, Rinse K. Weersma, Andre Franke, Daniel G. MacArthur, Chaim Jalas, Harry Sokol, Ramnik J. Xavier, Ann Pulver, Judy H. Cho, Dermot P.B. McGovern, Mark J. Daly

## Abstract

As part of a broader collaborative network of exome sequencing studies, we developed a jointly called data set of 5,685 Ashkenazi Jewish exomes. We make publicly available a resource of site and allele frequencies, which should serve as a reference for medical genetics in the Ashkenazim. We estimate that 30% of protein-coding alleles present in the Ashkenazi Jewish population at frequencies greater than 0.2% are significantly more frequent (mean 7.6-fold) than their maximum frequency observed in other reference populations. Arising via a well-described founder effect, this catalog of enriched alleles can contribute to differences in genetic risk and overall prevalence of diseases between populations. As validation we document 151 AJ enriched protein-altering alleles that overlap with “pathogenic” ClinVar alleles, including those that account for 10-100 fold differences in prevalence between AJ and non-AJ populations of some rare diseases including Gaucher disease (*GBA*, p.Asn409Ser, 8-fold enrichment); Canavan disease (*ASPA*, p.Glu285Ala, 12-fold enrichment); and Tay-Sachs disease (*HEXA*, c.1421+1G>C, 27-fold enrichment; p.Tyr427IlefsTer5, 12-fold enrichment). We next sought to use this catalog, of well-established relevance to Mendelian disease, to explore Crohn’s disease, a common disease with an estimated two to four-fold excess prevalence in AJ. We specifically evaluate whether strong acting rare alleles, enriched by the same founder-effect, contribute excess genetic risk to Crohn’s disease in AJ, and find that ten rare genetic risk factors in *NOD2* and *LRRK2* are strongly enriched in AJ, including several novel contributing alleles, show evidence of association to CD. Independently, we find that genomewide common variant risk defined by GWAS shows a strong difference between AJ and non-AJ European control population samples (0.97 s.d. higher, p<10^−16^). Taken together, the results suggest coordinated selection in AJ population for higher CD risk alleles in general. The results and approach illustrate the value of exome sequencing data in case-control studies along with reference data sets like ExAC to pinpoint genetic variation that contributes to variable disease predisposition across populations.

---

Recent advances in genome sequencing technologies are improving our understanding of the etiology of human diseases^1,2^. Similarly, epidemiological studies over the past century have improved our understanding of their global distribution^3–5^. To date, it remains unclear the extent to which genetics may play a role in population-based differences in prevalence and/or incidence. Efforts to increase inclusion of a broader representation of populations in genomic studies will likely improve our interpretation of these observed differences^6^. Here, we present a study on the relative contribution of DNA sequence variants to the risk and prevalence of Crohn’s disease (CD) and rare diseases in the Ashkenazi Jewish (AJ) population.

Genetic population isolates, defined as populations that start with a small group of founders and that may experience bottlenecks altering with periods of population growth^7^, have facilitated the mapping of alleles contributing to human disease predisposition^8–11^. Tight bottlenecks scatter the relative contributions of genes and as a consequence may make it easier to discover some disease-associated genes, although other genes, where the alleles may have been depleted from the isolated population, will be harder to detect^10^.

Genome-wide association studies (GWAS) and follow-up targeted sequencing efforts have been unusually successful in CD and have established a substantial role for low frequency variants across the more than 200 loci^2,12^ defined to date. In addition, the documented 2-4 fold enrichment of CD prevalence in the AJ population^13,14^, a population with an established founder effect, motivated the use of exome sequencing along with genome-wide array data to evaluate the degree to which bottleneck-enriched protein-altering alleles and unequivocally implicated common variants contribute an excess CD genetic risk to AJ, and as a consequence, to the documented increased prevalence of CD in AJ^13^ (Figure S1).

Additionally, founder effects, and possible selection, have made some rare diseases more prevalent in the Ashkenazi Jewish (AJ) population^15,16,17,18^, akin to the well-documented 40 rare diseases known as “Finnish heritage diseases”, which are much more common in Finland^9^ and whose difference in prevalence has largely been attributed to genetics. Despite the remarkable progress in mapping genes and alleles for some of these rare diseases, precise estimates of the risk-allele frequency and the carrier rate in the AJ population isolate have not yet been determined^16^. Through this study we provide a frequency resource of protein-coding alleles from over 2,000 non-CD AJ samples with low admixture fraction that will serve to improve interpretation of the carrier rate of rare disease risk alleles in the AJ population (Figure S1).

## Results

We generated a jointly called exome dataset consisting of 18,745 individuals from international IBD and non-IBD cohorts^1,19^. As the present study aimed to focus on variation observed in the AJ population in comparison to reference populations in ExAC^19,20^ (including non-Finnish Europeans (NFE), Latino (AMR), and African/African-American (AFR)) populations, we chose a model-based approach to estimate the ancestry of the study population using ADMIXTURE^21^.

To identify AJ individuals and estimate admixture fractions we included a set (n=21,066) of LD-pruned common (MAF>1%, see Supplementary Note for additional details) variants after filtering for genotype quality (GQ>20). The 18,745 samples were assigned to four groups (K=4) using ADMIXTURE. In one of the four groups, 3,522 samples had estimated ancestry fraction > 0.9, with the majority of the samples labelled as “AJ” by contributing study sites (Figure S1). Because we were interested in computing an enrichment statistic that would not be affected by possible admixture we obtained alternate (non-reference) allele frequency estimates by restricting the enrichment analysis to the 2,178 non-IBD Ashkenazi Jewish samples that passed QC and relatedness filtering and had AJ focused ancestry fraction > 0.9 (Figure S1).

To estimate parameters of enrichment in the AJ population, including proportion of enriched alleles and degree of enrichment, we used the observed alternate allele counts and total number of alleles to take into account uncertainty in estimated allele frequencies from AJ and NFE (n=33,370), AFR (n=5,203), and AMR (n=5,789) available from ExAC release 0.3 dataset (n_total_=60,706). We focused on protein-coding alleles with estimated allele frequency of at least 0.002 in AJ (n_alleles_=103,878), and applied a three group Bayesian mixture model, we refer to as the Population Isolate Enrichment Mixture Model (PIEMM) (see Supplementary Note), to classify the observed alleles into three groups: “depleted”, “similar”, or “enriched”. We estimate that 30% of the analyzed protein-coding alleles have a mean 7.6-fold increased frequency compared to reference populations with markedly different proportion of alleles belonging to the “enriched” group depending on variant annotation: 44% for predicted protein-truncating variants (PTV); 36% for predicted protein-altering variants (PRA); and 27% for synonymous variants (Figure 1, Figure S2, p < 10^−16^ across comparisons of PTV and PRA to synonymous variants, two-proportion test, Supplementary Note).

**Figure 1.**
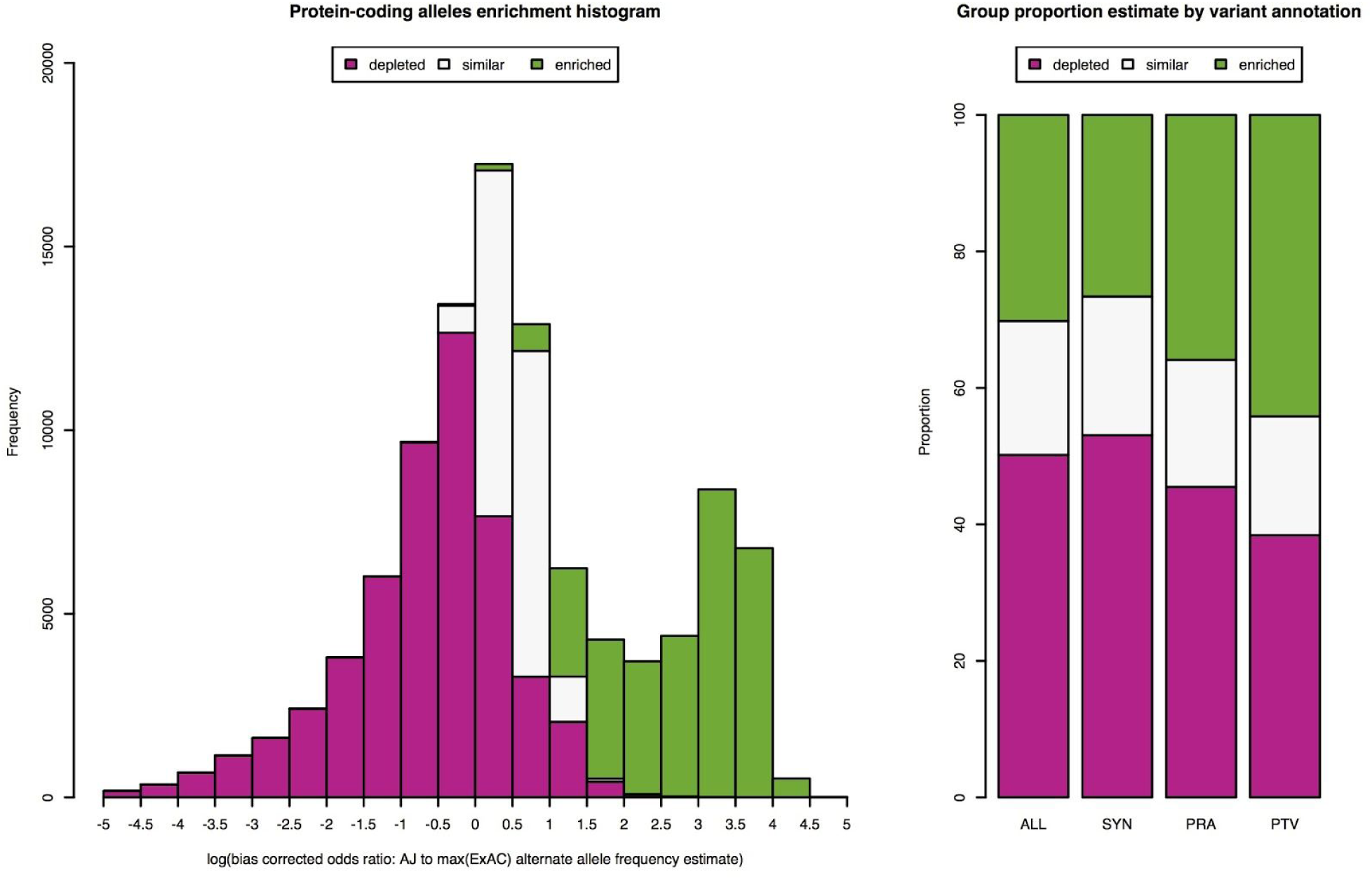
Enrichment of alleles discovered in AJ exome sequencing project. **A)** Density plot of estimated log enrichment statistic, defined as the log of the bias corrected odds ratio comparing the allele frequency in AJ population to the maximum allele frequency estimated from NFE, AFR, and AMR populations in ExAC. For each histogram bin we show a barplot of the expected number of alleles belonging to the three different groups we analyzed: 1) “depleted” (magenta); 2) “similar” (light gray); and 3) “enriched” (green). **B)** Bar plots of estimated proportion of alleles belonging to the three different groups we analyzed for all protein-coding (ALL), synonymous (SYN), protein-altering (PRA), and protein-truncating variants (PTV). An estimate of 30% of protein-coding alleles observed in AJ with a mean shift of 7.6-fold increase in allele frequency compared to other reference populations.

As validation of our approach to identify alleles that contribute to differences in genetic risk to disease, we intersected the list of protein-coding alleles identified in the AJ exome sequencing study with reported pathogenic and non-conflicted alleles (n=42,226) in ClinVar^22^ resulting in 151 alleles found both in ClinVar and with posterior probability greater than 0.5 of belonging to the enriched group (Table S1). In OMIM, 49 of the 151 alleles included documentation of a disease subject with AJ ancestry (Table 1). This set of enriched alleles includes 9/14 alleles described in the American College of Medical Genetics and Genomics 2008 screening guideline study for the AJ population^23^. In the setting of autosomal recessive disorders these differences in population allele frequencies may contribute to an order of squared enrichment difference in genetic risk and prevalence between populations (see Supplementary Note). For instance, a 12-fold enriched frameshift indel, p.Tyr427IlefsTer5, in *HEXA*, contributes a 144-fold enrichment in genetic risk in AJ to non-AJ population to Tay-Sach’s disease. This large cohort of adult Ashkenazi exome database further supports recent publications of founder mutations for rare pediatric disorders including: *FKTN* and Walker Warburg syndrome^24^; *CCDC65* and Primary ciliary dyskinesia^25^; *TMEM216* and Joubert syndrome^26^; *C11orf73* and Leukoencephalopathy^27^; *PEX2* and Zellweger syndrome^28^; *VPS11* and Hypomyelination and developmental delay^29^; and *BBS2* and Bardet-Biedl syndrome^30^. This resource will undoubtedly assist in prioritizing variants for gene discovery to further identify founder mutations in AJ (Table S2).

**Table 1.**
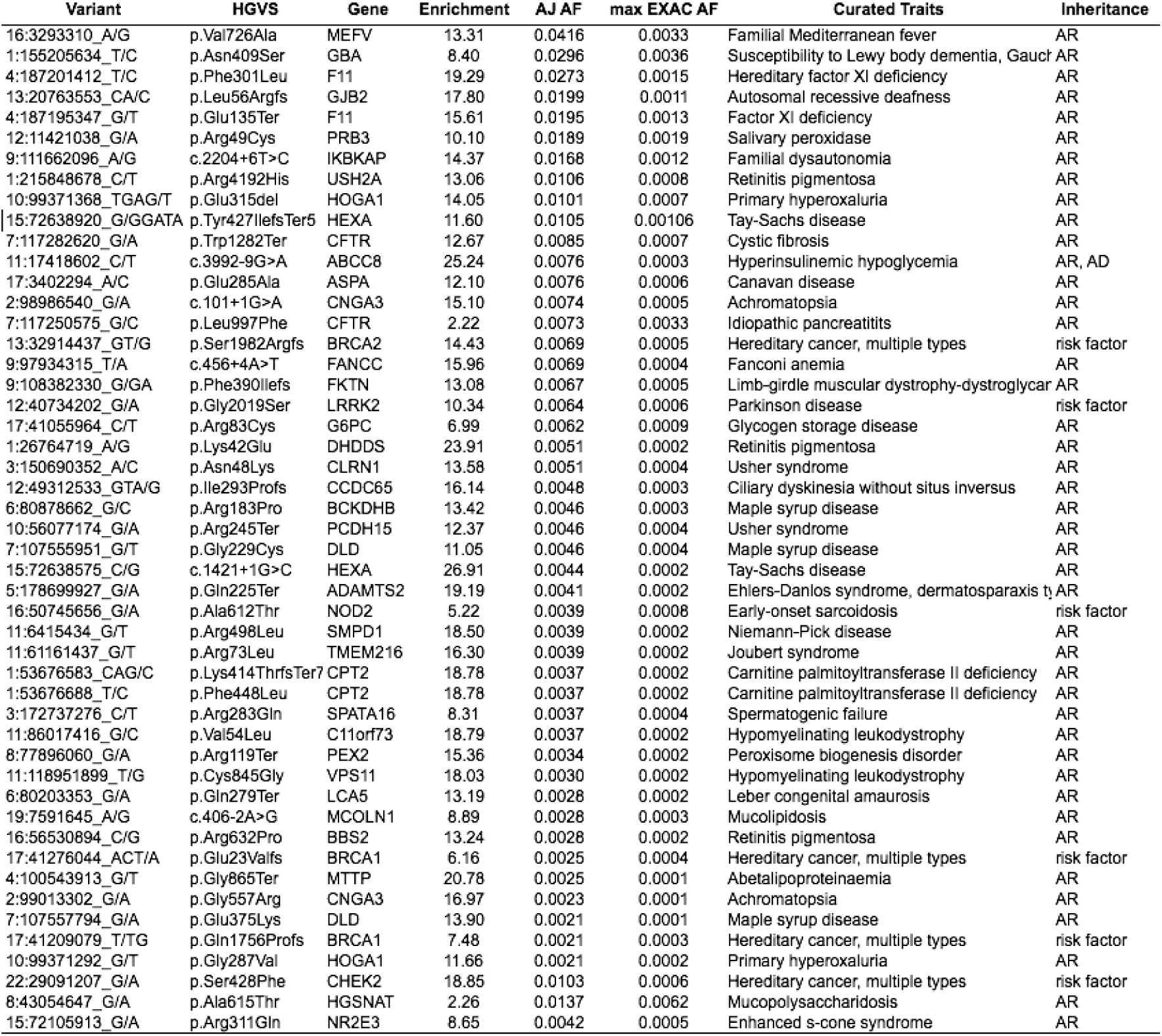
Forty-nine ClinVar “pathogenic” alleles enriched in AJ. HGVS and Gene is the allele nomenclature in ClinVAR and gene symbol. Enrichment corresponds to the comparison of allele frequency in AJ (AJ AF) to maximum frequency among three population groups (max EXAC AF): 1) NFE; 2) AMR; and 3) AFR. Curated trait is based on the trait description in the Online Mendelian Inheritance in Man (OMIM). Inheritance corresponds to the inheritance description in OMIM (AR: autosomal recessive, AD: autosomal dominant, risk factor: not specified genetic risk factor). Alleles are sorted in decreasing order by AJ AF.

To assess whether protein-coding alleles enriched in AJ population contribute to differences in CD genetic risk we performed case-control association analyses. Because individuals with partial AJ ancestry will still carry bottleneck-enriched alleles, here we included all samples with estimated AJ ancestry fraction of at least 0.4 (Figure S2), which resulted in a dataset of 4,899 AJ samples (1,855 Crohn’s disease and 3,044 non-IBD). To improve ability to detect association we performed a meta-analysis with CD and non-IBD case-control exome sequencing data from two separate ancestry groups: 1) non-Finnish European (NFE) (2,296 CD and 2,770 non-IBD); and 2) Finnish (FINN) (210 CD and 9,930 non-IBD samples) from a separate callset described in a previous publication^31^ for a total of 4,361 CD samples and 15,744 non-IBD samples.

Study-specific association analysis was performed with Firth bias-corrected logistic regression test^32,33^ and four principal components as covariates using the software package EPACTS^34^ (Figure S4). We combined association statistics in a meta-analysis framework using the Bayesian models in Band et al^35^. We used the correlated effects model, obtained a Bayes factor (BF) by comparing it with the null model where all the prior weight is on an effect size of zero, reported p-value approximation using the BF as a test statistic, and assessed whether heterogeneity of effects exist across studies for downstream QC (see Supplementary Note). We separately assessed CD observed vs. expected associations for enriched protein-altering (pra) and synonymous (syn) alleles in protein-coding genes in CD implicated GWAS loci (n_gwas,pra_=413; n_gwas,syn_=231), and outside implicated GWAS loci (n_non-gwas,pra_=14,961; n_non-gwas,syn_=7,858, Figure 2).

**Figure 2.**
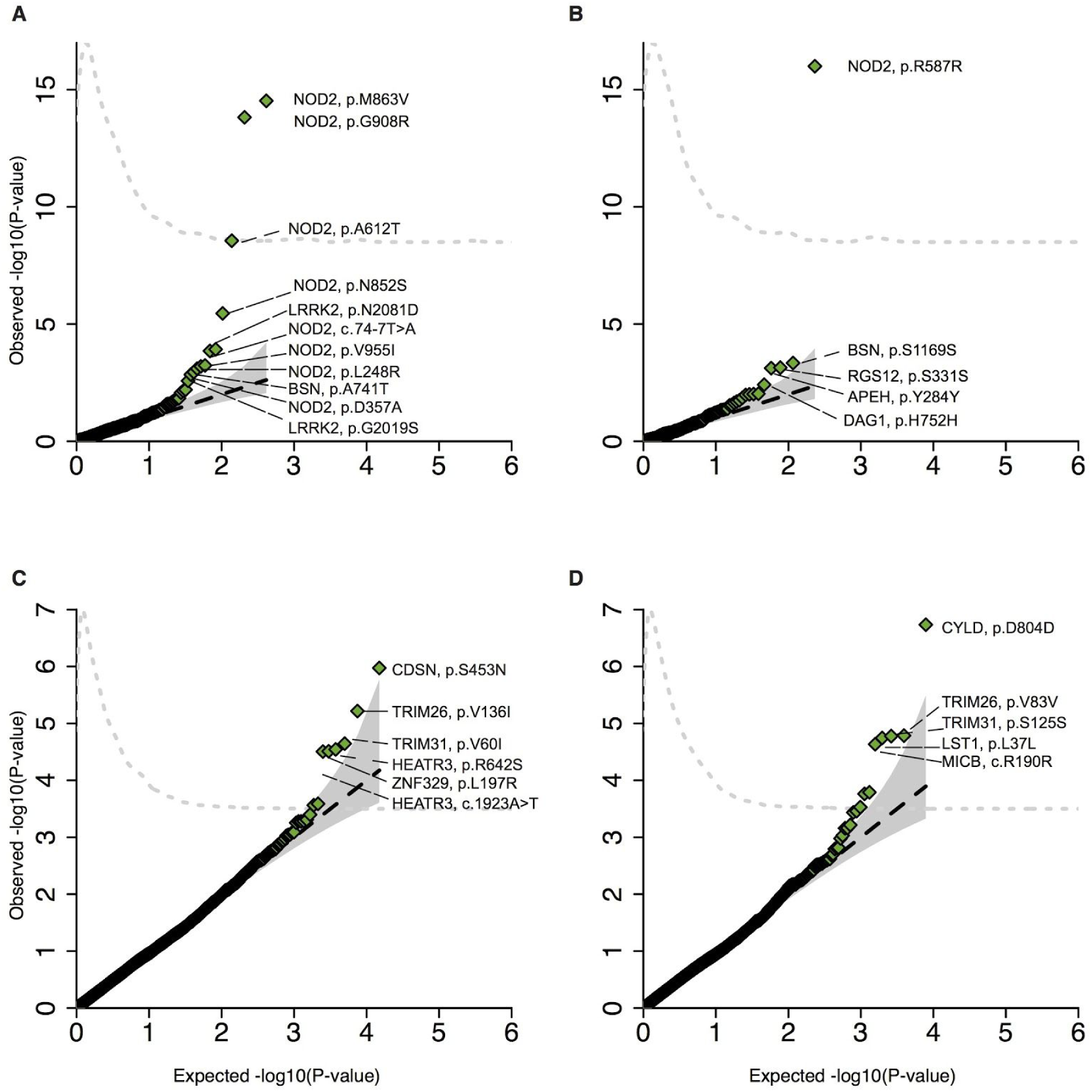
Q-Q plots of enriched alleles. Q-Q plots of Crohn’s disease association for: **A**) AJ enriched protein-altering (protein-truncating and missense) and **B**) synonymous alleles in GWAS regions; and AJ enriched **C**) protein-altering and **D**) synonymous alleles outside of GWAS regions. For each Q-Q plot variants with a corresponding p-value less than or equal to a threshold where expected number of false discoveries is equal to one are annotated. The black dashed line is y = x, and the grey shapes show 95% confidence interval under the null. The gray dashed line represents the observed density of -log10 p-values.

We identified ten AJ enriched CD risk alleles (p<0.005): the previously published risk haplotypes in *LRRK2* and *NOD2* (*LRRK2*: p.N2081D; *NOD2*: p.N852S, p.G908R, p.M863V+fs1007insC)^36,2^, in addition to newly implicated alleles (*NOD2*: p.A612T, p=2.8x10^−9^; c.74-7T>A, p=1.4×10^−4^; p.L248R, p=6.4×10^−4^; p.D357A, p=0.0011; *LRRK2*: p.G2019S, p=0.0014, a Parkinson’s disease risk allele^37^). To assess whether the new *NOD2* enriched alleles are conditionally independent of the previously established associated *NOD2* alleles we performed conditional haplotype association analysis in PLINK and Bayesian model averaging^38^ for variable selection, both of which suggested independent effects for all alleles (Figure S5, Table S3).

Despite the functional relationship between *LRRK2* and *NOD2*^39^, we do not observe deviation from additivity between *LRRK2* and *NOD2* (p=0.273). Deviation from additivity has been reported for p.fs1007insC, p.G908R, and p.R702W in *NOD2*^40,41^. We assessed whether any independent evidence of deviation from additivity exists for the newly associated single nucleotide substitutions. In agreement with previous reports, deviation from additivity existed with estimated genotype odds ratios of 1.84 for heterozygous and 7.39 for compound heterozygous/homozygous genotypes (p=0.0038, analysis of deviance, ANOVA), and found significant evidence of deviation from additivity for the newly reported alleles (p=0.00357, odds ratio = 7.53). We found no evidence of deviation from additivity for the associated protein-altering alleles in *LRRK2* (p=0.418).

Given the presence of genetic variants in *NOD2* and *LRRK2* that contribute to differences in genetic risk in AJ population, we next asked whether unequivocally established common variant loci associations may also contribute to differences in genetic risk. We performed polygenic risk score (PRS) analysis using reported effect size estimates from 124 CD alleles including those reported in a previously published study^12^ and four variants in *IL23R* from a recent fine-mapping study^42^, and excluding variants in *NOD2* and *LRRK2*. We observed an elevated PRS for AJ compared to non-Jewish controls (0.97 s.d. higher, p<10^−16^; Figure 3A; number of non-AJ controls=35,007; number of AJ controls=454). We observed a similar trend for the CD samples (0.54 s.d. higher; p<10^−16^; Figure 3B; number of non-AJ CD cases=20,652; number of AJ CD cases=1,938).

**Figure 3.**
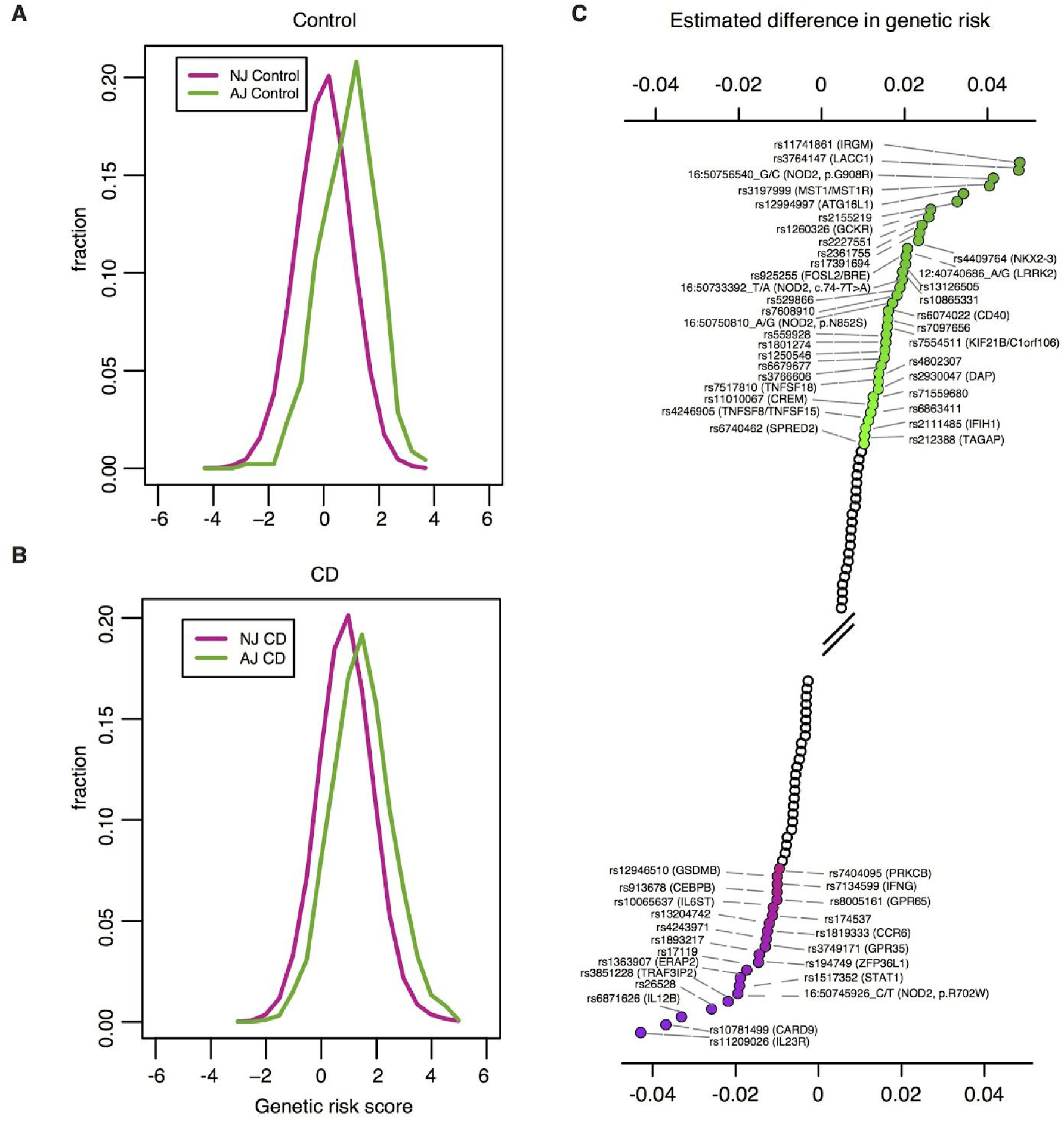
AJ individuals have higher CD polygenic risk score than NJ controls. NJ: non-Jewish; AJ: Ashkenazi Jewish; CD: Crohn’s disease; PRS: polygenic risk score. **A**) Density plot of CD polygenic risk scores in 454 AJ (green) and 35,007 NJ (purple) controls. AJ controls have elevated CD polygenic risk score that NJ controls (0.97 s.d. higher, p<10^−16^). **B**) Density plot of CD polygenic risk scores in 1,938 AJ (green) and 20,652 NJ CD (purple) cases (0.54 s.d. higher, p<10^−16^). For both density plots the scores have been scaled to NJ controls, thus resulting in an NJ control PRS density of mean equal to 0 and variance equal to 1 (see Online methods). **C**) Ranked (decreasing order) CD associated variants by estimated contribution to the differences in genetic risk between AJ and NJ. Associated variants with estimated contribution greater than or equal to 0.01, computed as 2*log(odds ratio)*(AJ frequency - NJ frequency), assuming additive effects on the log scale, are highlighted in green. Associated variants with estimated contribution less than or equal to −0.01 are highlighted in purple. Forward slashes represent a break in variants highlighted.

To quantify the relative contribution of CD-implicated alleles to the difference in genetic risk between AJ and non-AJ populations we estimated the expected PRS value of an individual and expected difference in PRS between two populations by simply using summary statistics including the frequency of the minor allele in the two populations and the corresponding odds ratio (Supplementary note, Figures S6-S8).

We applied the approach to all CD implicated alleles and observed that variants in GWAS loci annotated as *IRGM*, *LACC1*, *NOD2*, *MST1*, *ATG16L1*, *GCKR, NKX2-3*, and *LRRK2*^12^ contribute substantially (>0.01) to the increased genetic risk observed in AJ. It is possibly relevant that variants contributing to increased risk in AJ correspond to autophagy/intracellular defense genes (*IRGM, ATG16L1, LRRK2*) and those contributing to increased risk in non-AJ correspond to anti-fungal/Th17/ILC3 genes^43^ (*IL23R*, *IL12B*, *CARD9*, *TRAF3IP2*, *IL6ST*, *CEBPB*; Figure 3C).

Two factors impact our ability to obtain precise estimates of the contribution of genetic risk factors to the documented differences in disease prevalence between populations. First, contemporaneous epidemiological data are largely unattainable. This confounds our ability to obtain any informative estimates due to documented variability in the occurrence of CD over time^3,44^. Second, there has been substantial uncertainty in reported CD prevalence estimates^45,46^. Here, to interpret the impact of shifts in genetic risk score on differences in prevalence, we used the logit risk model^41^ and evaluated a new estimate of disease probability, p_new_, assuming an initial disease probability, p_0_, and multiple values for the differences in genetic risk. Assuming log-additive effects, and a log-risk model, we estimate that the observed differences in genetic risk between the AJ and non-AJ populations may contribute an expected 1.5-fold increase in disease prevalence in a population with environmental risk factors corresponding to AJ and baseline genetic risk corresponding to non-AJ populations (Figure S6-S8). To address the extent to which non-additive effects in *NOD2* may impact the observed prevalence we assumed shared heterozygous and compound heterozygous/homozygous odds ratios of 1.84 and 7.39, respectively. We estimate a 1.6% increase in difference in prevalence attributed to the deviation from additivity, suggesting a small effect on differences in population prevalence (Supplementary Note).

## Discussion

By drawing on data from 5,685 Ashkenazi Jewish exomes we provide a systematic analysis of AJ enriched protein-coding alleles, which may contribute to differences in genetic risk to CD as well as numerous other rare diseases. We identified protein-altering alleles in *NOD2* and *LRRK2* that are conditionally independent and contribute to the excess burden of CD in AJ. We found evidence that common variant risk defined by GWAS shows a strong elevated difference between AJ and non-AJ European population samples, independent of *NOD2* and *LRRK2*^47^, suggesting a coordinated selection in AJ for higher CD risk alleles^48–50^.

We made a couple of unique observations by studying CD in the AJ population. First, studying recently bottlenecked populations enables powerful discovery of genetic variation that markedly differ in frequency and as a consequence contributes to differences in genetic risk across population groups. Second, *NOD2* and published common variant associations contribute substantially to the genetic risk of CD, making it more difficult to identify other genes whose ancestral alleles failed to pass through the bottleneck, consistent with predictions from Zuk et al^10^.

Finally, we provide an exome frequency resource of protein-coding alleles in AJ along with estimates of enrichment. Our approach and this resource will likely catalyze our understanding of the medical relevance of enriched alleles in population isolates.

## Author contributions

M.A.R, D.P.B.M, M.J.D participated in the study design. M.A.R, J.K., M.K., and M.J.D analyzed whole exome data. M.A.R, H.H, T.H, and B.A analyzed the SNP chip data. B.M.N, A.G, D.G, B.G, I.P, G.A, N.B, A.P.L, E.S, N.P, Ben Weisburd, K.J.K, E.V.M, B.P, L.B, P.S, J.C, Graham Heap, T.A, V.P, A.W.S, S.T, Dan Turner, P.S, M.F, K.K, M.P, Aarno Palotie, S.R.B, D.G.M, R.H.D, Mark S. Silverberg, J.D.R, R.K.W, A.F, H.S, R.J.X, A.P, J.H.C, D.P.B.M provided reagents, methods, and tools for analysis. C.S. managed the project. C.J and J.H.C provided detailed analysis of rare diseases in AJ. All authors commented on the manuscript. J.D.R, Mark J. Daly, S.R.B, R.H.D, Mark S. Silverberg, J.H.C, and D.P.B.M are members of the NIDDK IBD Genetics consortium. Manuel A. Rivas, J.K., B.A, and Mark J Daly wrote the manuscript.

## Acknowledgments

M.J.D. is supported by grants from the following: the National Institute of Diabetes and Digestive and Kidney Disease (NIDDK) and the National Human Genome Research Institute (NHGRI; DK043351, DK064869 and HG005923); the Crohns and Colitis Foundation (3765); the Leona M. & Harry B. Helmsley Charitable Trust (2015PG-IBD001); the Stanley Center; and Amgen (2013583217). R.J.X. is supported by grants from Amgen (2013583217) and CCFA (3765). J.D.R. is funded by grants from NIDDK (DK064869 and DK062432). G.A. is supported by NIH R01 grant AG042188. N.B. is supported by NIH grants AG618381, AG021654, AG038072, and the Glenn Center for the Biology of Human Aging. A.F. and B.P. are supported by the DFG (Deutsche Forschungsgemeinschaft) Cluster of Excellence “Inflammation at Interfaces” and the DFG grant FR 2821/6-1. I.P. is supported by the Leona M. and Harry B. Helmsley Charitable Trust and New York Crohn’s Foundation. IBD Research at Cedars-Sinai is supported by grant PO1DK046763, and the Cedars-Sinai F. Widjaja Foundation Inflammatory Bowel and Immunobiology Research Institute Research Funds. D.P.B.M. is supported by DK062413, AI067068 and U54DE023789-01; grant 305479 from the European Union; and The Leona M. and Harry B. Helmsley Charitable Trust and the Crohn’s and Colitis Foundation of America. S.R.B. is support by an NIH U01 grant (DK062431). J.H.C. is supported by grants from NIH (U01 DK062429, U01 DK062422, R01 DK092235, SUCCESS), and the Sanford J. Grossman Charitable Trust. H.S. is supported by Equipe ATIP–Avenir 2012 grant and INSERM–ITMO SP 2013. E.V.M. is supported by an NIH F13 award (AI122592-01A1). Researchers at UCL are supported by the Charles Wolfson Foundation Trust. RKW is supported by a VIDI grant (016.136.308) from the Netherlands Organization for Scientific Research (NWO). R.H.D. is supported by NIH grant U01 DK062420 and the Inflammatory Bowel Disease Genetic Research Chair at the University of Pittsburgh. We thank Dr. Jonathan Bloom for proposed edits and comments on the manuscript. We thank the Broad IT team for assistance with the IBD exomes browser.

## S1. Materials and methods

### Initial variant call set

We generated a jointly called dataset consisting of 18,745 individuals from international IBD and non-IBD cohorts. Sequencing of these samples was done at Broad Institute.

### Ethics statement

All patients and control subjects provided informed consent. Recruitment protocols and consent forms were approved by Institutional Review Boards at each participating institutions (Protocol Title: The Broad Institute Study of Inflammatory Bowel Disease Genetics; Protocol Number: 2013P002634). All DNA samples and data in this study were denominalized.

### Cohort descriptions

For all cohorts, CD was diagnosed according to accepted clinical, endoscopic, radiological and histological findings.

### Target selection

G4L WES is a project specific product. It combines the Human WES (Standard Coverage) product with an Infinium Genome-Wide Association Study (GWAS) array. In addition to the array adding to the genomics data, it also acts as a concordance QC, linking 14 SNPs to the exome data. The processing of the exome includes Sample prep (Illumina Nextera), hybrid capture (Illumina Rapid Capture Enrichment - 37Mb target), sequencing (Illumina, HiSeq machines, 150bp paired reads), Identification QC check, and data storage (5 years). Our hybrid selection libraries typically meet or exceed 85% of targets at 20x, comparable to ~60x mean coverage. The array consists of a 24-sample Infinium array with ~245,000 fixed genome-wide markers, designed by the Broad. On average our genotyping call rates typically exceed 98%.

### Pre-processing

The sequence reads are first mapped using BWA MEM^51^ to the GRCh37 reference to produce a file in SAM/BAM format sorted by coordinate. Duplicate reads are marked – these reads are not informative and are not used as additional evidence for or against a putative variant. Next, local realignment is performed around indels. This identifies the most consistent placement of the reads relative to potential indels in order to clean up artifacts introduced in the original mapping step. Finally, base quality scores are recalibrated in order to produce more accurate per-base estimates of error emitted by the sequencing machines.

### Variant Discovery

Once the data has been pre-processed as described above, it is put through the variant discovery process, i.e. the identification of sites where the data displays variation relative to the reference genome, and calculation of genotypes for each sample at that site. The variant discovery process is decomposed into separate steps: variant calling (performed per-sample), joint genotyping (performed per-cohort) and variant filtering (also performed per-cohort). The first two steps are designed to maximize sensitivity, while the filtering step aims to deliver a level of specificity that can be customized for each project.

Variant calling is done by running Genome Analysis Toolkit’s (GATK) HaplotypeCaller in GVCF mode on each sample’s BAM file(s) to create single-sample gVCFs. If there are more than a few hundred samples, batches of ~200 gVCFs are merged hierarchically into a single gVCF to make the next step more tractable. Joint genotyping is then performed on the gVCFs of all available samples together in order to create a set of raw SNP and indel calls. Finally, variant recalibration is performed in order to assign a well-calibrated probability to each variant call in a raw call set, and to apply filters that produce a subset of calls with the desired balance of specificity and sensitivity as described in Rivas et al. (2016)^52^. Samples with >= 10% contamination are excluded from call sets. Exome samples with less than 40% of targets at 20X coverage are excluded.

### Variant annotation

Variant annotation was performed using the Variant Effect Predictor (VEP) [cite PMID: 20562413] version 83 with Gencode v19 on GRCh37. Loss-of-function (LoF) variants were annotated using LOFTEE (Loss-Of-Function Transcript Effect Estimator, available at <u>https://github.com/konradjk/loftee</u>), a plugin to VEP. LOFTEE considers all stop-gained, splice-disrupting, and frameshift variants, and filters out many known false-positive modes, such as variants near the end of transcripts and in non-canonical splice sites, as described in the code documentation.

### Identification of Finnish samples

Finnish CD patients were recruited from Helsinki University Hospital and described in more detail previously^53,54^. We used the same exome sequencing dataset described in Rivas et al.^31^ We applied additional PC correction in the Finnish identified individuals to remove individuals with membership of Finnish sub-isolate (Northern Finland) and excluded based on PC2 >= 0.015 (853 excluded, 826 controls, 27 IBD). We recalculated PCs and included the first four PCs in the association analysis.

### Ancestry estimation and quality control

As the present study aimed to focus on variation observed in Ashkenazi Jewish (AJ) population in comparison to reference populations in ExAC^55^ including (non-Finnish Europeans (NFE), Latino (AMR), and African/African-American (AFR)) we chose a model-based approach to estimate the ancestry of the study population using ADMIXTURE^21^. To identify AJ individuals and estimate admixture proportions we included a set (n=21,066) of LD-pruned common variants (MAF>1%) variants after filtering for genotype quality (GQ>20). We selected 50 window size in SNPs, 5 SNPs to shift the window at each step, and the variance inflation factor (VIF) threshold equal to 2. The 18,745 samples were assigned to four groups (K=4) using ADMIXTURE. In one of the four groups, 3,522 samples had estimated ancestry fraction > 0.9, with the majority of the samples labelled as “AJ” by contributing study sites, belonging to the group with high probability (Figure S1B).

Prior to enrichment and association analysis, 81 samples (of total 18,745) were also filtered due to possible contamination (heterozygous/homozygous ratio < 1), excess of singletons (n>2000), deletion/insertion ratio (>1.5) and mean genotype quality (<40). 275 samples were excluded for relatedness (pi>0.35 cut-off). Genotypes with low genotype quality (<20) were filtered, in addition to variants with low call rate (<80%) and allele balance deviating from 70:30 ratio for greater than 40% of heterozygous samples if at least 7 heterozygous samples were identified.

As we were interested in computing an enrichment statistic that would not be affected by possible admixture, we obtained alternate allele frequency estimates by restricting the enrichment analysis to the 2,178 non-IBD Ashkenazi Jewish samples that passed QC and relatedness filtering and had AJ focused ancestry fraction > 0.9 (Figure S1). Principal Component Analysis (PCA) was done in each ancestry group using the 21,066 variants. Sample QC was done using the Hail software while PCA, differential missingness, allele balance and sample relatedness analysis was done using PLINK^56^. Hail is an open-source software framework for scalably and flexibly analyzing large-scale genetic data sets (https://github.com/broadinstitute/hail).

### Estimating fold-enrichment in AJ population compared to reference populations in ExAC

#### Statistical methods: Population Isolate Enrichment Mixture Model (PIEMM)

To estimate which alleles are enriched in AJ compared to alleles in reference population groups in ExAC we developed a statistical method we refer to as the **P**opulation **I**solate **E**nrichment **M**ixture **M**odel (**PIEMM**). This model estimates the proportion of observed alleles in the population isolate that are enriched or depleted as well as the shift in distribution of those alleles relative to the reference populations.

Using the number of alternate and reference alleles observed in AJ non-IBD samples and in the population (NFE, AFR or AMR) with the highest frequency frequency from ExAC we compute a bias corrected log odds ratio estimate, 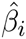, and its standard error, 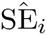, for odds of the alternate allele as

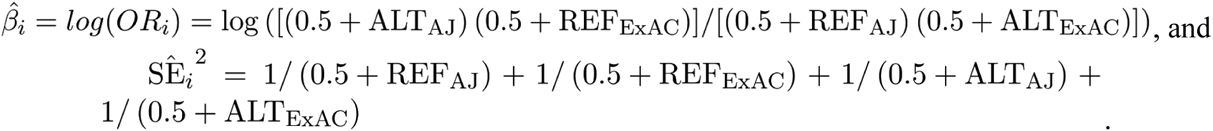

These formulas can be understood in a Bayesian framework to be mean posterior estimates of *β*_*i*_ and SE_*i*_ based on the ExAC and observed AJ allele frequencies and a prior distribution of Beta(½, ½). This is the Jeffreys prior distribution for the parameter of a binomial distribution.

We use these summary statistics to estimate:

i. which proportion of alleles observed in AJ belong to each of three groups: (0) ‘similar’ - AJ allele has similar frequency to reference populations, (1) ‘enriched’ - AJ allele has shifted increased frequency to reference populations, (2) ‘depleted’ - AJ allele has shifted decreased frequency to reference populations; and
ii. the extent to which enriched and depleted groups are shifted in frequency from the reference populations.

#### Details

Let 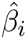 and 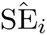 be the log odds ratio and standard error estimates of protein-coding allele *i* in AJ with respect to the reference population with the highest allele frequency. Let *γ*_*i*_ be a variable that is either 0, 1, or 2. Our PIEMM is the following mixture model:

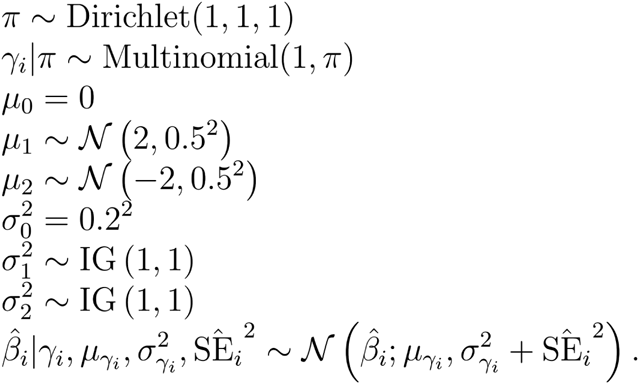

#### Motivation of parameters and distribution

The group membership of each AJ protein-coding allele is unknown in advance. As a result the proportion *π* of the alleles belonging to the similar, enriched, or the depleted group (characterized by an unknown shift in mean *μ* and variance *σ*^2^) needs to be estimated. Our prior for *π* is the uniform distribution on the 2d simplex (also known as the ^Dirichlet(1,1,1)^ distribution) so as to not favor *a priori* any particular value of *π*.

The prior for the variance parameter *σ*^2^ is the inverse gamma distribution with hyperparameter values *α* = 1 and *β* = 1 for the enriched and the depleted group. This distribution is relatively flat between 0.3 and 1, and thus covers well the region where we expect the variance parameter to reside. The prior for the variance parameter *σ*^2^ for the ‘similar’ group is fixed to 0.2^2^, thus including alleles in the similar group where small deviation of enrichment is observed. The prior shift hyperparameter values for the ‘depleted’ and the ‘enriched’ group are −2 and +2, respectively, indicating separation in shift.

#### MCMC algorithm

We use a Gibbs sampler, an approximation algorithm, to approximate the posterior distribution of the parameters of the PIEMM. Superscripts for the variables denote their value after the corresponding iteration. Let *l* be the group index whenever it is not explicitly given.

1. Initialize 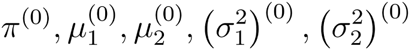, and 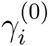 for all *i*.
2. Repeat for *t* = 1, 2,… *n*_burn_ + *n*_iter_
  a. For *i* = 1, 2,…, *n*_alleles_, generate 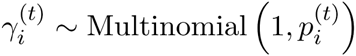 where 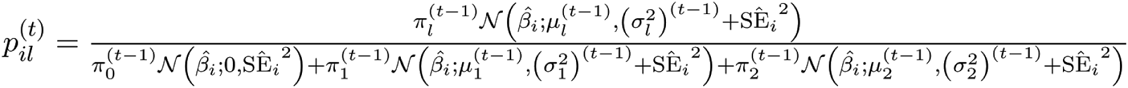, for *l* = 0, 1, 2.
  b. Generate *π*^(*t*)^ ~ Dirichlet (1 + *n*_0_, 1 + *n*_1_, 1 + *n*_2_), where 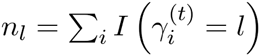, for *l* = 0, 1, 2.
  c. Update: 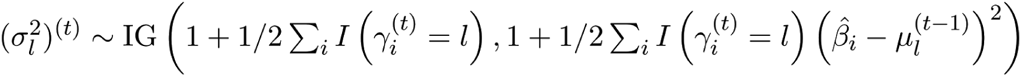, and 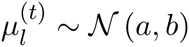, where

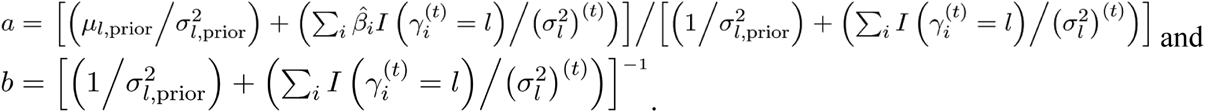

We run the algorithm for *n*_burn_ + *n*_iter_ = 1000 iterations and discard the first *n*_burn_ = 100 iterations from the analysis. Figure S3 shows a typical example of a plot used to evaluate the performance of the algorithm across all the iterations.

To estimate allele enrichment in AJ compared to reference populations we used 2,178 non-IBD Ashkenazi Jewish samples, after sample and relatedness QC.

We calculated alternate allele frequencies for the Ashkenazi Jewish population and used allele frequency information for NFE (n=33,370), AFR (n=5,203), and AMR (n=5,789) available from ExAC release 0.3 dataset (n_total_=60,706) and focused on alleles where allele frequency information was available for AJ and the reference populations. For the enrichment plot we focused on alleles with estimated frequency of at least 0.002 in AJ (n_alleles_=103,878) and with alleles observed with an estimated frequency of at least .0001 in the reference populations with depth of coverage of at least 20X in at least 80% of the samples in ExAC.

#### Overlap of enriched alleles with ClinVar

We harmonized the XML and TXT releases of the ClinVar database (April 11, 2016 data release) ^22^ into a single tab-delimited text file using scripts that we have released publicly (<u>https://github.com/macarthur-lab/clinvar</u>). Briefly, we normalized variants using a Python implementation of vt normalize^57^ and de-duplicated to yield a dataset unique on chromosome, position, reference, and alternate allele. A variant was considered ‘pathogenic’ if it had at least one assertion of either Pathogenic or Likely Pathogenic for any phenotype. A variant was considered ‘conflicted’ if it had at least one assertion of Pathogenic or Likely Pathogenic, and at least one assertion of Benign or Likely Benign, each for any phenotype. By these criteria, ClinVar contained n=42,226 identified as pathogenic and non-conflicted. Intersecting with our dataset revealed that 151 belonged to the AJ enriched group with high probability (>.5).

#### Assessing Crohn’s disease association of protein-coding variation that may contribute to difference in disease prevalence in AJ

We focused Crohn’s disease association analysis of protein-coding variant to alleles that may account for difference in disease prevalence in AJ population to reference populations. To do so we focused on alleles with high probability of belonging to the enriched group. We included all samples with ADMIXTURE estimated AJ ancestry fraction of at least 0.4 (we excluded any samples that had alternative ancestry fraction of at least .4 in any other group). Samples with Ulcerative Colitis (n=700), unspecified and Indeterminate Colitis (n=86) were excluded from subsequent analysis. This resulted in a dataset of 4,899 AJ samples (1,855 Crohn’s disease and 3,044 non-IBD).

Study-specific association analysis was performed with Firth bias-corrected logistic regression test^32,33^ and four principal components as covariates using the software package EPACTS version 3.2.6 ^34^. Minimum minor allele count (≥1) and variant call rate (≥0.8) filters were used.

For meta-analysis we combined association statistics using the Bayesian models and frequentist properties proposed in Band et al^35^, which is a normal approximation to the logistic regression likelihood suggested by Wakefield^58^. As the authors of Band et al. indicate one way of thinking about the approach is that it uses the study-wise estimated log-odds ratio (beta) and its standard error as summary statistics of the data. For each model of association we assume a prior on the log odds ratio which is normally distributed around zero with a standard deviation of 0.2). By changing the prior on the covariance (or correlation) in effect sizes between studies we can formally compare models where: 1) the effects are independent across studies, and 2) the effects are correlated equally between studies. For each model we can obtain a Bayes factor (BF) for association by comparing it with the null model where all the prior weight is on an effect size of zero. We report p-value approximation using the Bayes factor as a statistic for model 2 where the effects are correlated between studies.

Association statistics were combined based on association analysis across three study groups: 1) AJ (1,855 CD and 3,044 non-IBD samples); 2) NFE (2,296 CD and 2,770 non-IBD); and 3) Finnish (FINN) (210 CD and 9,930 non-IBD samples) for a total of 4,361 CD samples and 15,744 non-IBD samples.

#### Conditional haplotype based testing and variable selection for *NOD2* alleles

In the conditional haplotype based testing (--chap) analysis we used PLINK v1.08p^56^ and set a minimum haplotype frequency of .001 (--mhf). We used PLINKSEQ (<u>https://atgu.mgh.harvard.edu/plinkseq/</u>), an open-source C/C++ library for working with human genetic variation data, and the Python bindings implemented in pyPLINKSEQ to perform Bayesian Model Averaging (BMA). We applied BMA^38^ using the R package ‘BMA’ (<u>https://cran.r-project.org/web/packages/BMA/BMA.pdf</u>).

#### Polygenic risk scores

The polygenic risk scores were calculated for the international inflammatory bowel diseases consortium European samples. Details of these samples including the QC procedures were described in previous publications ^42^. We used reported effect size estimates from 124 CD alleles including those reported in a previously published study^12^ and four variants in *IL23R* from a recent fine-mapping study^42^, and excluding variants in *NOD2* and *LRRK2*. We used 454 AJ controls; 1,938 AJ CD; 35,007 non-Jewish controls and 20,652 non-Jewish CD samples. Polygenic risk scores were calculated using array genotype data as the sum of the log odds ratio of the variants associated with CD. Scores for missing genotypes were replaced by the imputed expected value using PLINK ^56^. Variants in *NOD2* and *LRRK2* were excluded from the analysis to assess whether polygenic signal was independent.

Let PRS_*i*_ be the polygenic risk score of individual *i*, assuming additive effects on the log-odds scale then

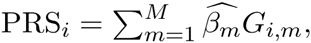

where 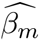 denotes the estimated log odds ratio for variant *m* and *G*_*i,m*_ denotes the genotype dosage of individual *i* for variant *m*.

In the setting where effects are non-additive, i.e. a genotype-specific effect model, then 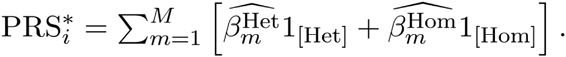.

For now, we consider the additive scenario, and later we return to the setting where non-additive effects exist, which is relevant for quantifying the contribution of *NOD2* alleles to differences in genetic risk between two populations.

The estimated expected PRS value for an individual in population *j* is

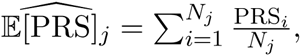

where *N*_*j*_ is the number of individuals sampled in population *j*. Substituting equation for *PRS*_*i*_ and rearranging terms simplifies the equation as a function of variant frequency:

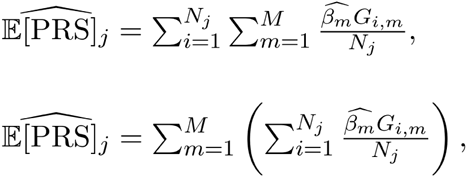

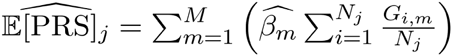, where 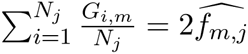, and 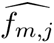 denotes the frequency of variant *m* in population *j*. Thus, the estimated expected PRS value of an individual in population *j* is 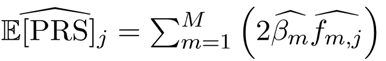.

Assume that we are interested in the expected difference in PRS between two individuals, say from population 1 being AJ and population 2 being NFE. Also, assume that the effect size of variant *m* is shared across both populations. Then, using the estimated expected PRS value we define estimated expected difference in PRS as the difference in estimated expected PRS value between two populations:

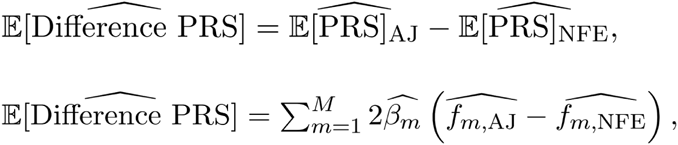

which can be used to get an estimated contribution of a variant *m* to the difference in polygenic risk score between two populations,

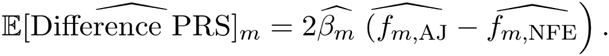

To rank variants according to their relative contribution to differences in genetic risk we included the *NOD2* and *LRRK2* alleles, used the list of estimated effect size from the published studies ^12,42^, and estimates from this study (Supplementary Table 3).

If we replace PRS* for PRS

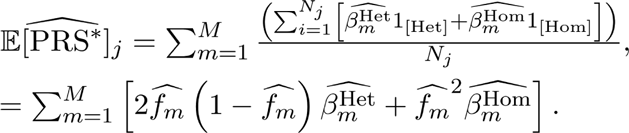

Then, the estimated expected difference in PRS* when non-additive effects exist is

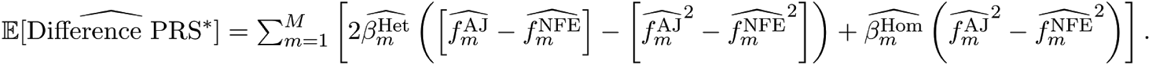

#### Estimating fold difference in prevalence for a population with shift in expected genetic risk

Assuming log-additive effects in the logit risk model the disease probability for an individual is given as *p* = (1 + exp (−*η*))^−1^, where *η* tends towards a normal distribution with parameters 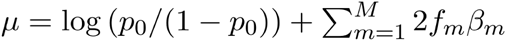 and 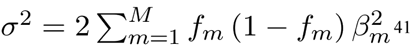. Here *p*_0_ refers to a baseline disease probability.

We can see that *μ* may be expressed in terms of the expected polygenic risk score, i.e. 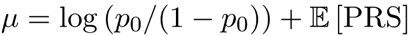. In the setting where 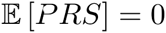 then 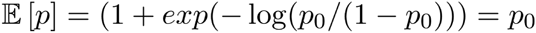.

To evaluate the impact of a shift in the expected value of polygenic risk score to the expected value of *μ* we can express the shift as:

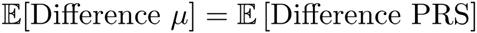. We can compute new values of *p* for new values of *μ* to obtain a fold-increase in prevalence for a population that has undergone such a shift.

We see that this requires a value to be chosen for *p*_0_ and that log(*p*_0_/(1−*p*_0_)) can be represented as a baseline risk score value *β*_0_. To get an estimate of the absolute prevalence of CD in the AJ population, we must choose a baseline *β*_0_, where *p*_0_ represents the expected prevalence with zero non-baseline alleles in the population^41^, to which we add a contribution from multiple non-baseline alleles to calculate: 1) an individual’s probability of disease, or 2) the expected prevalence of the disease in the population.

Once we have chosen a value for *β*_0_, we can calculate the ratio of expected prevalence as follows. First, use the means (*μ*_AJ_ and *μ*_NAJ_) and variances (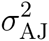 and 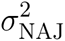) of risk scores as calculated above to calculate the probability density function of the disease prevalence. In the case of the AJ population, we have

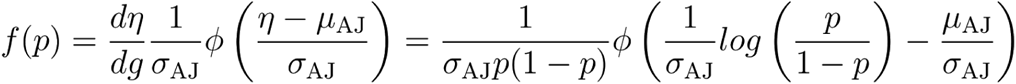

where *η* is the risk score associated with prevalence *p,g* is the link function, so *p* = *g*(*η*) = (1 + *e*^(−*η*)^)^−1^, and *ϕ* is the standard normal density function.

Next, we integrate to get 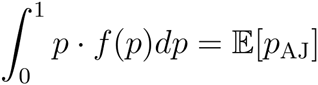. Finally, we can calculate 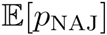 in a similar way, and divide the expected prevalence in the AJ population by that in the non-AJ population to get the prevalence ratio, 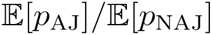.

The value of *β*_0_ = −20.5 was chosen in order to obtain a prevalence in the non-AJ population of ~0.5%. At this value of *β*_0_, the ratio of prevalence in the AJ population to that in the non-AJ population was estimated to be 1.5 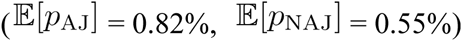.

For different choices of *β*_0_, however, this ratio may vary, as the relationship between probability of disease and risk score is non-linear. Figure S6 shows how the values of the disease prevalence and their ratio vary as *β*_0_ is changed. We see that the ratio values range from 1.46 to 1.52 for different values of *β*_0_ with a range of baseline prevalence of .001 to .01 - the range of prevalence estimates for Crohn’s disease^3,46,59^.

To further understand the effect that choosing a logit based model had on the results, a comparison of the standard logit and probit models was done using the values inferred from the logit model. No full scale probit modelling was done in this analysis, so the values found with the probit model represent only a close approximation of the expected results.

In the logit model for population analysis, we may assume that individual risk scores are chosen from a normal distribution 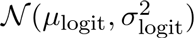 where *μ*_logit_ and *σ*_logit_ represent the mean and standard deviation of the risk scores as defined above. From here, we may calculate the probability density function of probit model risk scores *η*_probit_ based on that of logit model risk scores *η*_logit_ as

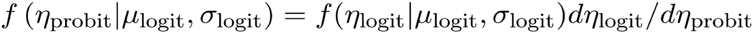

and use this to calculate *μ*_*probit*_ and *σ*_*probit*_, the estimated mean and standard deviation of the risk scores in the probit model. Using these values, we obtain a probability distribution for the frequency of disease in the populations using the probit model.

While the logit model yielded a prevalence ratio of 1.506, the probit estimation yielded a prevalence ratio of 1.5136, with similar expected prevalence values 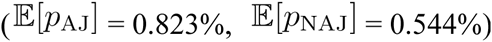. These values demonstrate that individual logit and probit analyses would likely give extremely similar results for values of interest. The complete probability densities under the logit and probit models can be seen in Figure S7.

Further, it is interesting to compare the relationship between values of risk scores in the two models. For values of risk scores between −1 and 1 in the logit model, the relationship to those in the probit model is highly linear, with a formula of *η*_probit_ = 0.6223 · *η*_logit_, with r^2^ = 1.0000. This formula may be used to impute single values in one model or the other assuming that the estimated total risk score is otherwise close to zero, and the imputed value is low. It is worth noting, however, that this does not work for all values of *η*_logit_, as the relationship between risk score in the logit and probit models deviates from this simple linear model when the risk score values are large.

#### Difference in prevalence between AJ and NFE attributed to implicated variants

The difference in prevalence due to multiple alleles can be computed as

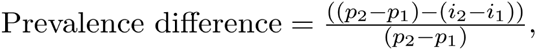

where *p*_*j*_ denotes the disease prevalence in population *j* and *i*_*j*_ denotes the disease prevalence without the risk factors in population *j*, which according to Moonesinghe et al. ^60^ is

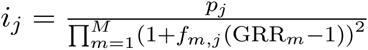

where GRR_*m*_ denotes the genotype relative risk for variant *m*.

We model the CD prevalence accounted for by CD associated enriched protein-altering alleles separately in both AJ and non-AJ European and determine the amount that CD prevalence would be reduced if this variant were absent from each population. Two-to-four fold difference in prevalence has been documented to exist between these groups^13^.We use a standard log-additive effect model and assume an odds ratio and uncertainty as estimated in the meta-analysis for each variant in both populations.

To estimate the difference in prevalence between two populations attributed to genetic risk factors when non-additive effects exist

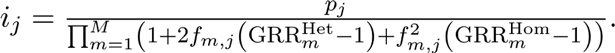

## S2. Online resources

IBD Exomes Browser: <u>http://ibd.broadinstitute.org</u>

Clinvar table: <u>https://github.com/macarthur-lab/clinvar/blob/master/output/clinvar.tsv</u>

ExAC sites VCF: <u>ftp://ftp.broadinstitute.org/pub/ExAC_release/release0.3/</u>

---

**Supplementary Figure 1.**
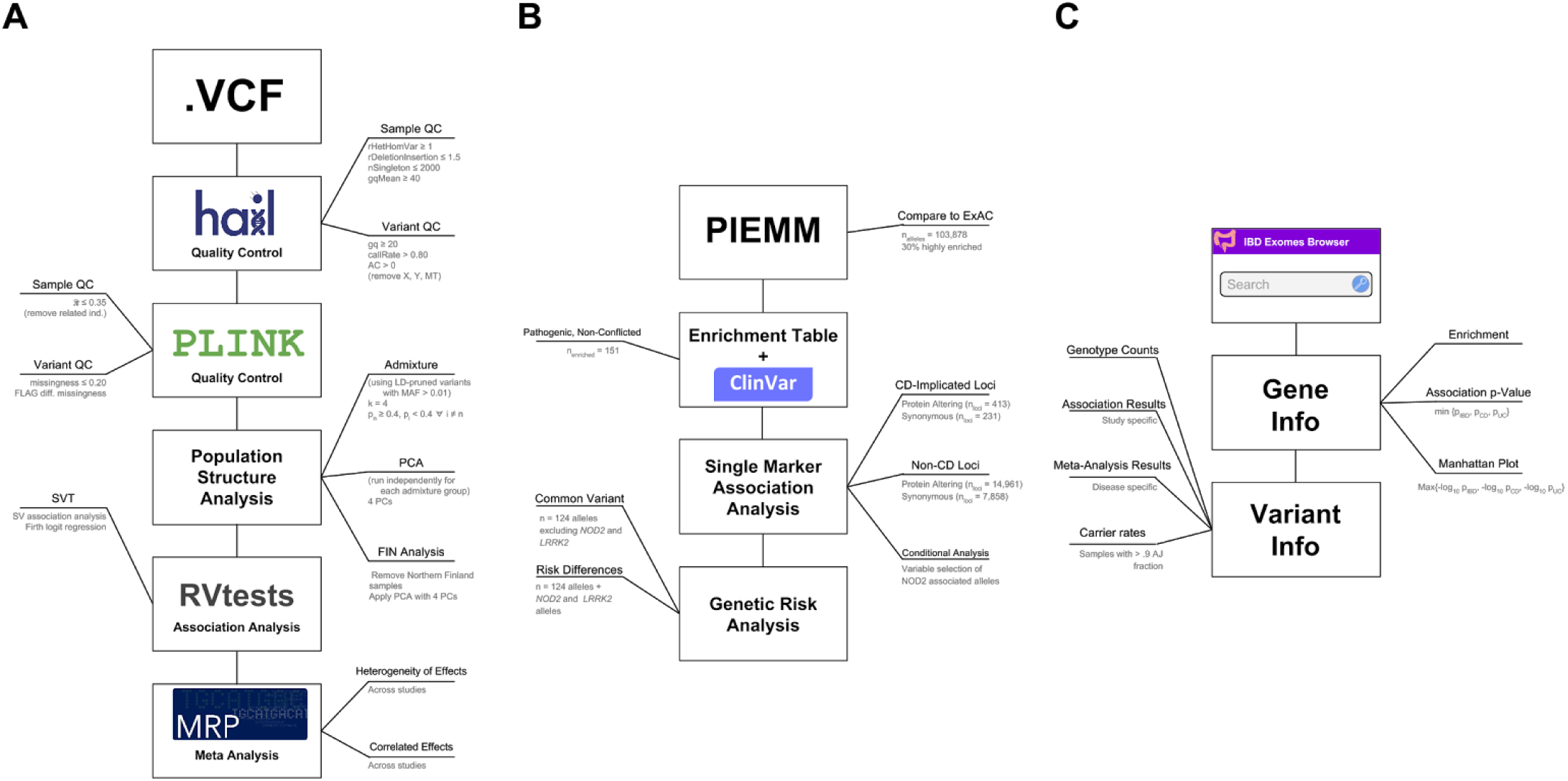
Analysis workflow diagram. **A)** Quality control, population structure, and association analysis workflow. **B)** AJ specific analysis workflow. **C)** Results and summary statistics are uploaded to the IBD browser hosted at: <u>http://ibd.broadinstitute.org</u> - a website that contains a gene and variant search engine, a “Gene Info” landing page that contains a manhattan plot and additional summary statistics, and a detailed “Variant Info” page that contains additional information about the alleles identified in our exome sequencing studies.

**Supplementary Figure 2.**
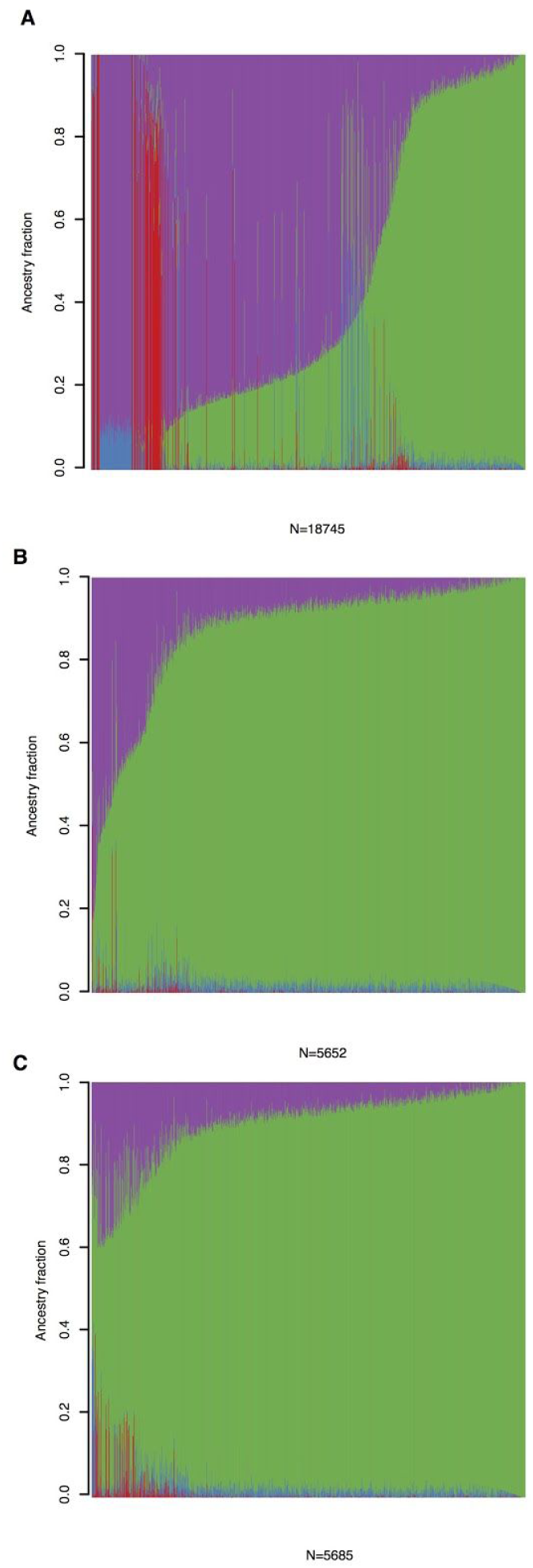
Admixture plots. **A)** Admixture plot, with 4 groups (K=4), for all samples exome sequenced. On the y-axis shown are ancestry mixture fractions for each of the samples (on x-axis) exome sequenced. **B)** Admixture plot for samples with self-reported Ashkenazi Jewish ancestry. **C)** Admixture plot for final samples used in the AJ analysis. All plots are ordered by ancestry fraction mostly loading for AJ samples (green). Group mostly loading for NFE samples is highlighted in magenta, East-Asian in red and African-American in blue.

**Supplementary Figure 3.**
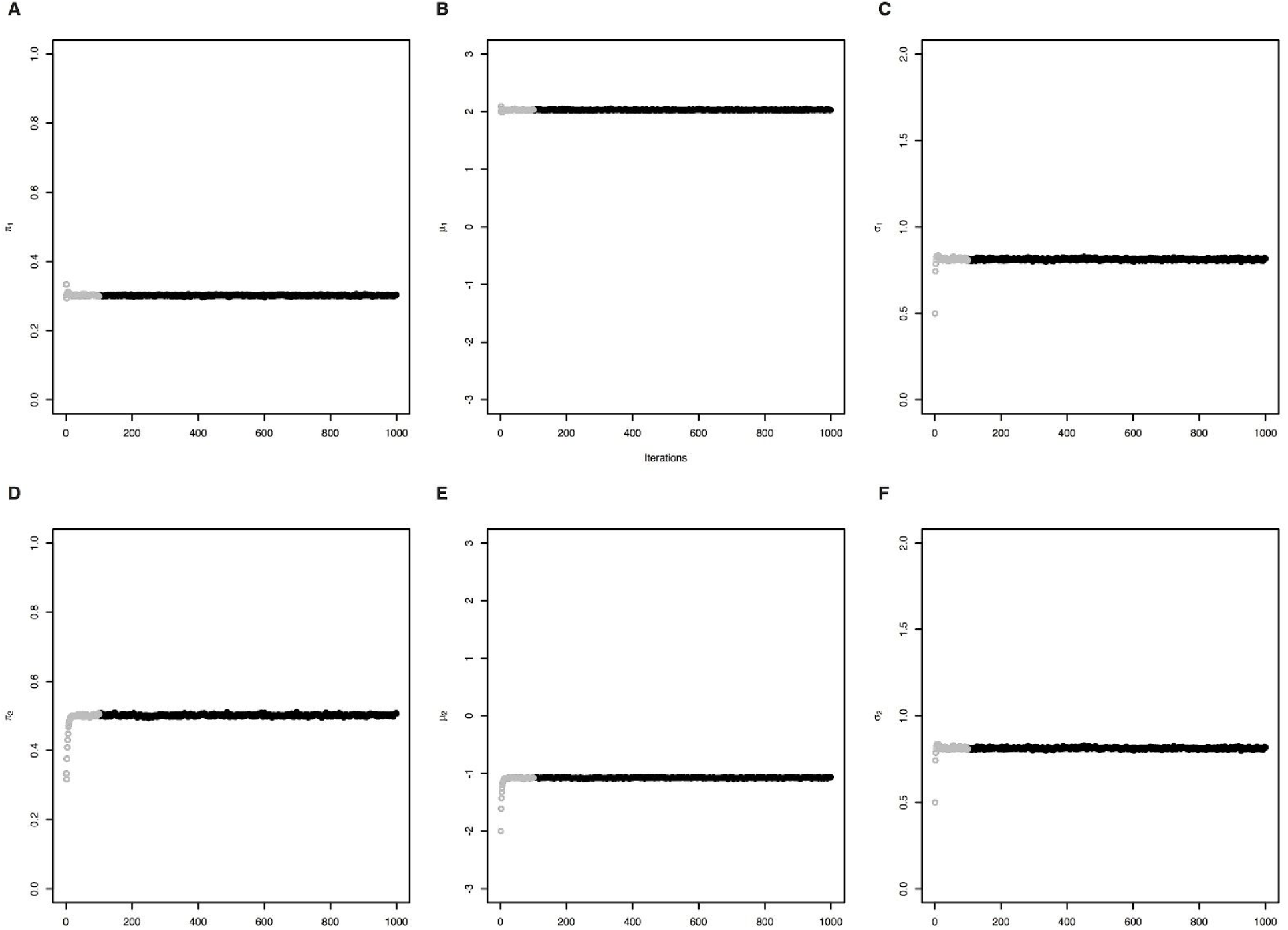
Parameter estimates for MCMC chain. When applying MCMC algorithms to a data set it is customary to show the performance of the algorithm across all stages of the experiment (including the burn-in). We show that the PIEMM algorithm generates stable proportion estimates for the two groups: “enriched” **(A-C)** and “depleted” **(D-F)** for parameters: *π* **(A, D)**, the proportion of alleles belonging to the group; *μ* **(B, E)**, the shift of the distribution belonging to the group; and *σ* **(C, F)**, the standard deviation of the distribution belonging to the corresponding group. For each group we demonstrate the parameter estimate during the burn-in (100 iterations, gray circles) and non burn-in (900 iterations, black circles) stage of the experiment used to obtain the parameter estimates reported in the manuscript.

**Supplementary Figure 4.**
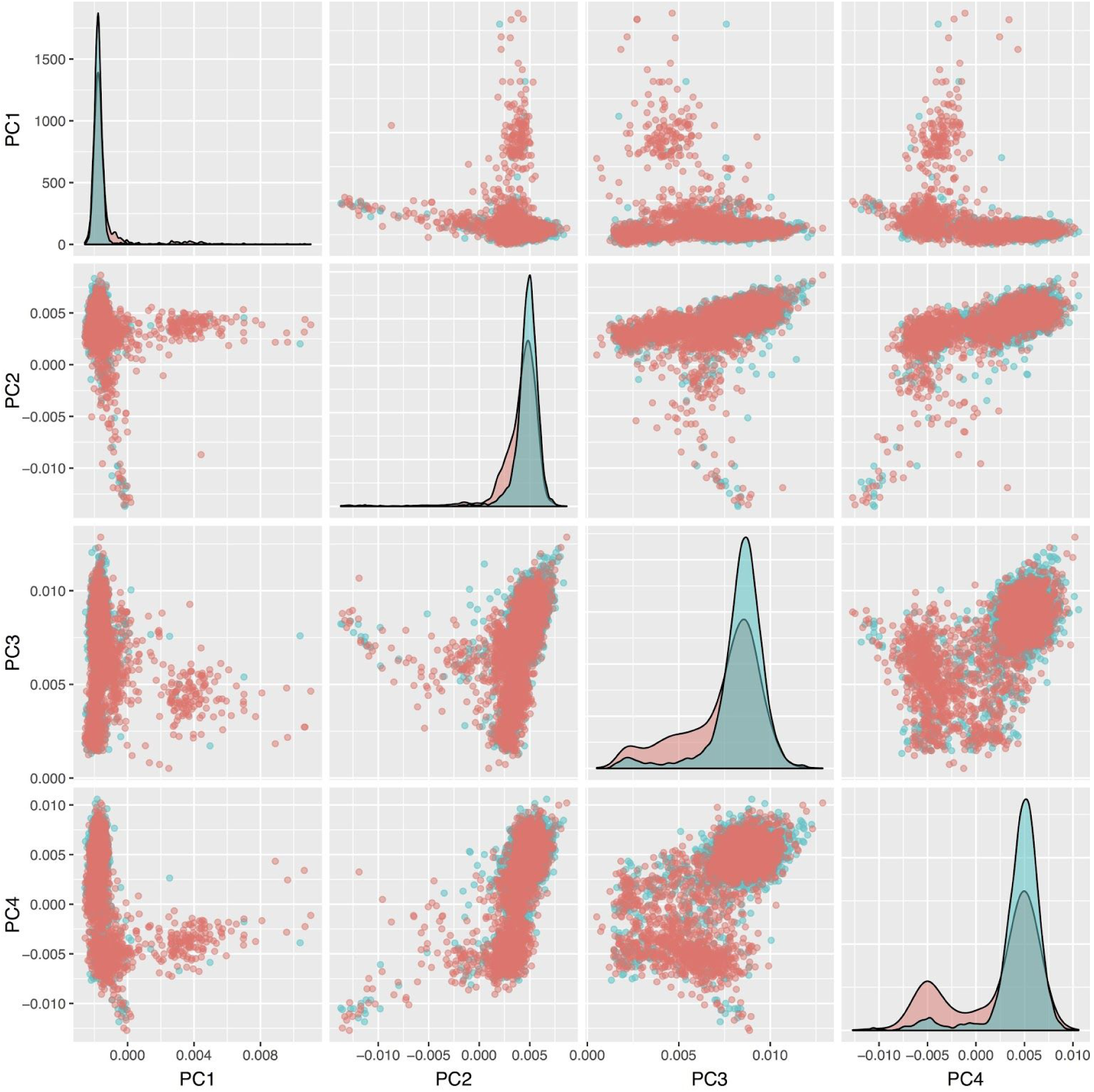
Principal components plot for 5,685 AJ individuals. Density plots for each of the PCs is shown (diagonal) separately for IBD cases (red) and controls (blue). Pairwise scatter plots are shown for PC1-PC4 separately for IBD cases and controls.

**Supplementary Figure 5.**
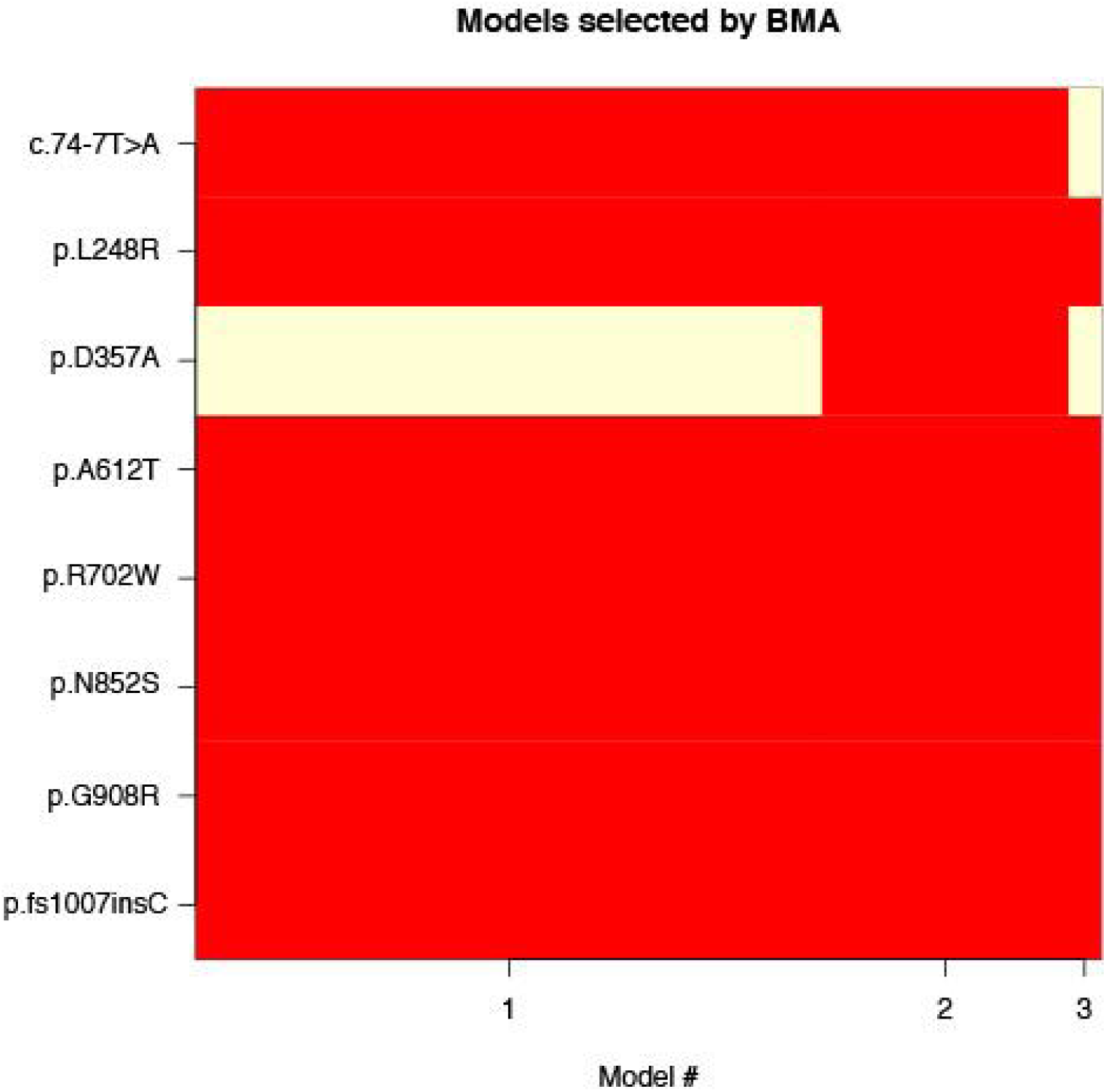
Variable selection using Bayesian model averaging (BMA). Eight protein-altering alleles with evidence of association and their corresponding membership for the models with high probability (> 0.01) after applying BMA, which accounts for the model uncertainty inherent in the variable selection problem by averaging over the best models in the model class according to approximate posterior model probability. Model #1 corresponds to the model where 7/8 alleles have a non-zero effect (p.D357A is not included) with approximate posterior model probability of .692. Model #2 corresponds to the model where 8/8 alleles have a non-zero effect with approximate posterior model probability of .273. Model # 3 corresponds to the model where 6/8 alleles have a non-zero effect (p.D357A and c.74-7T>A are not included) with approximate posterior model probability of .035.

**Supplementary Figure 6.**
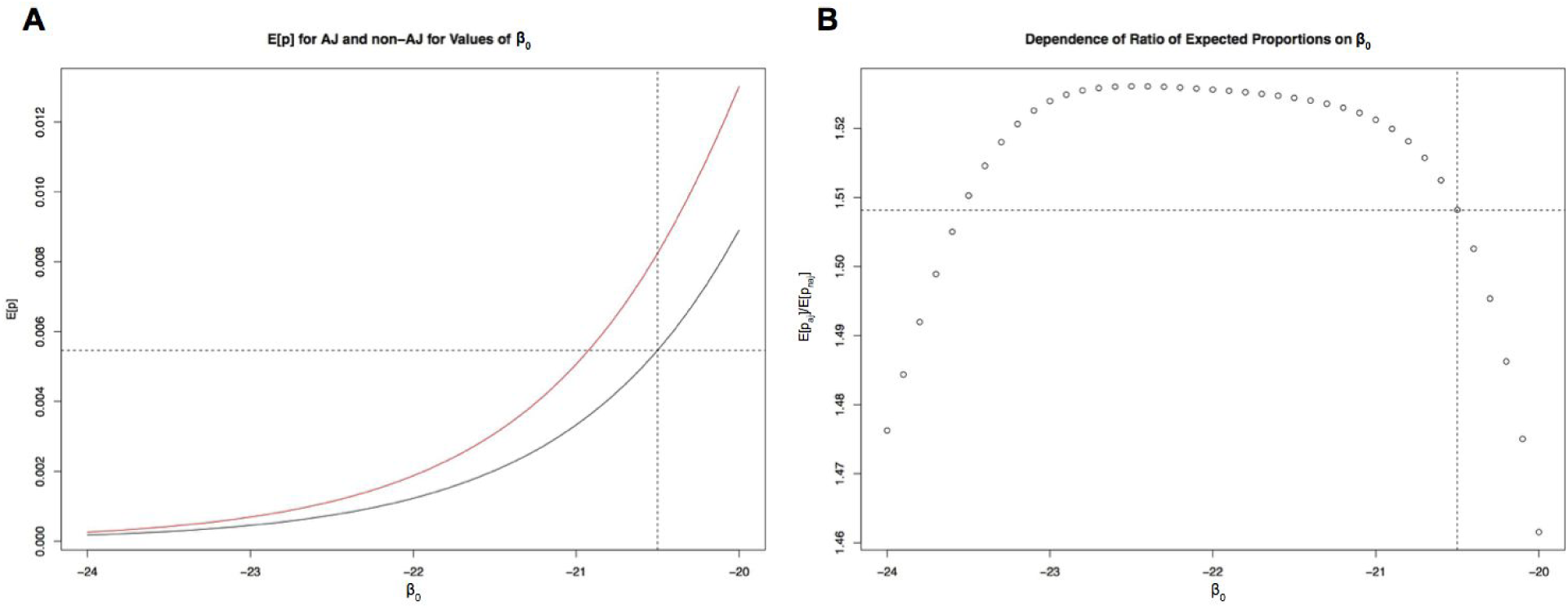
Dependence of population prevalence on *β*_0_. **A)** Expected values of disease prevalence in non-AJ (black) and AJ (red) populations. The value of *β*_0_ = −20.5 was chosen to approximate the prevalence of disease in the non-AJ population at around 0.005. **B)** Relationship between expected prevalence ratio and choice of *β*_0_. The chosen value of *β*_0_ = −20.5 corresponds to a ratio of 1.506.

**Supplementary Figure 7.**
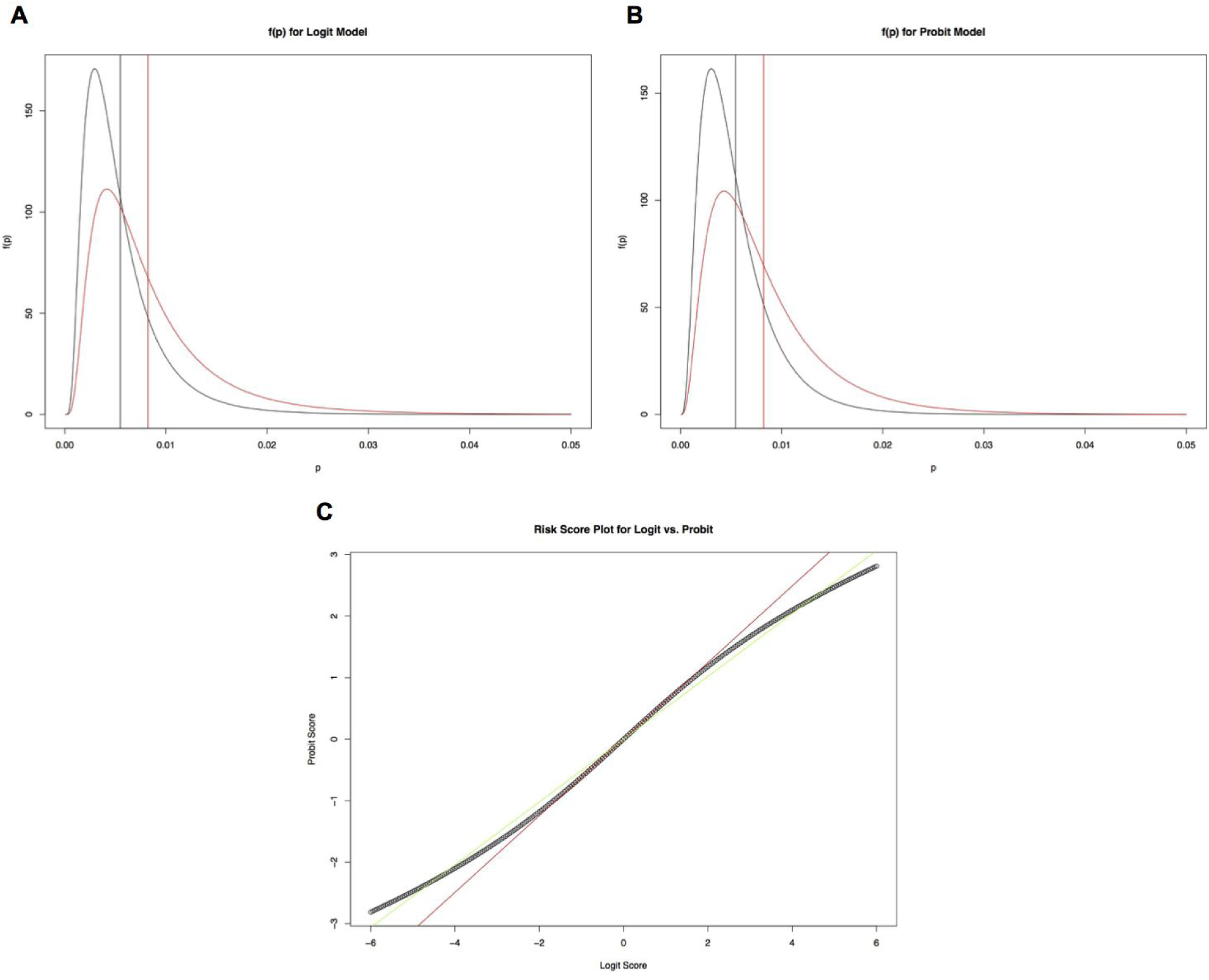
Probit and logit model analysis. **A)** Probability distribution of prevalence in non-AJ (black) and AJ (red) populations given the calculated mean and variance of risk score given by the logit model. **B)** Equivalent probability distributions given the calculated mean and variance of the risk score in the probit model. **C**) A comparison of risk score values in logit and probit models. The green line corresponds to a linear fit for the entire shown region, and the red line corresponds to a linear fit for the linear range around zero.

**Supplementary Figure 8.**
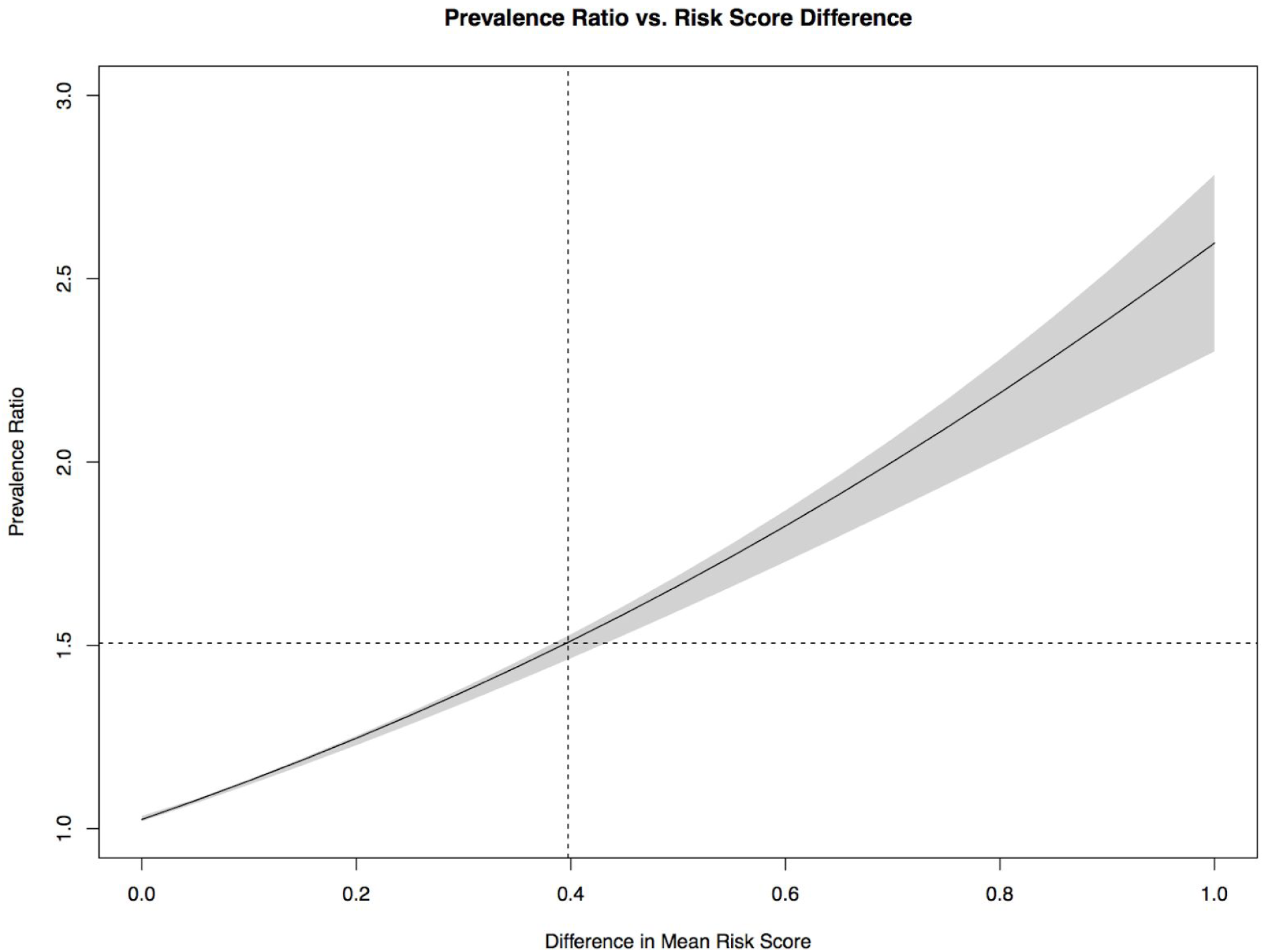
The relationship between expected differences in genetic risk score and expected fold difference in disease prevalence. Vertical line represents the estimated expected difference in genetic risk between AJ and non-AJ European population (0.397). For varying levels of expected difference in genetic risk we compute the expected fold-difference in prevalence. The shaded region marks the range of prevalence ratios obtained by varying *β*_0_ in a region of reasonable estimates (−24 to −20).

**Supplementary Table 3.** Conditional haplotype-based testing in *NOD2*. HGVS nomenclature for each allele (HGVS) and corresponding p-values are shown for independent effects given the haplotype formed by residual variants.

| Variant | HGVS | P-value |
| --- | --- | --- |
| 16:50763778:G:GC | p.fs1007insC | 6.50E-24 |
| 16:50745926:C:T | p.R702W | 2.18E-05 |
| 16:50756540:G:C | p.G908R | 1.52E-20 |
| 16:50750810:A:G | p.N852S | 1.78E-05 |
| 16:50745656:G:A | p.A612T | 3.98E-08 |
| 16:50733392:T:A | c.74-7T>A | 2.07E-05 |
| 16:50744565:T:G | p.L248R | 0.00198 |
| 16:50744892:A:C | p.D357A | 0.0016 |

